# Detoxification of endogenous serine prevents cell lysis upon glucose depletion in bacteria

**DOI:** 10.1101/343921

**Authors:** Michelle A. Kriner, Arvind R. Subramaniam

**Affiliations:** Basic Sciences Division and Computational Biology Section of the Public Health Sciences Division, Fred Hutchinson Cancer Research Center, Seattle, WA 98109, USA

**Author notes:** Corresponding author (ARS).

## Abstract

The amino acid serine, despite its diverse metabolic roles, can become toxic when present in excess. Indeed, many bacteria rapidly deaminate exogenously supplied serine into pyruvate and ammonia, even at the expense of biomass production. Here we report a surprising case in which endogenously produced serine must be detoxified in order for the bacterium *Escherichia coli* to survive. Specifically, we show that *E. coli* cells lacking the *sdaCB* operon, which encodes a serine transporter and a serine deaminase, lyse upon glucose depletion when serine is absent from the growth medium. Lysis can be prevented by omission of glycine or by inhibition of the glycine cleavage system, suggesting that activation of glycine catabolism upon glucose depletion causes a transient increase in intracellular serine levels. Heterologous expression of the serine transporter SdaC is sufficient to prevent lysis, indicating a dominant role for serine export, rather than deamination, in mitigating serine toxicity. Since lysis can be modulated by altering alanine availability, we further propose that mis-incorporation of serine instead of alanine into peptidoglycan crosslinks is the cause of lysis. Together, our results reveal that SdaC-mediated detoxification of intracellularly produced serine plays a protective role during sudden shifts in nutrient availability in bacteria.

**Author summary:** The amino acid serine is a building block used to make many types of macromolecules, yet bacteria actively degrade serine that is provided in growth media. Serine degradation is thought to prevent toxic serine accumulation, but the biological role of this process is not fully understood. We observed that cells lacking the *sdaCB* operon, which encodes a serine transporter and an enzyme that converts serine to pyruvate, suddenly lyse upon depletion of glucose from the growth medium. This surprising phenotype occurs only in media lacking serine, suggesting that *sdaCB* is required to detoxify intracellularly produced serine. Expression of the serine transporter SdaC is sufficient to prevent lysis, providing the first evidence that serine export can be an essential function of this protein. Our results reveal that sudden shifts in nutrient availability can increase the intracellular concentration of useful metabolites to toxic levels and suggest that increasing intracellular serine levels by manipulating SdaC activity may be a possible antimicrobial strategy.

## Introduction

The amino acid serine is a centrally important biomolecule, not only as a substrate for protein synthesis but also as a precursor of nucleotides, redox molecules, phospholipids and other amino acids^1,2^. While the biochemical reactions involved in serine metabolism are well known, our understanding of how cells regulate fluxes and intracellular concentrations of serine-associated metabolites remains incomplete. Recent revelations that serine biosynthesis genes are amplified or up-regulated in many cancers^3–6^ and required for host colonization by bacterial pathogens^7,8^ has led to renewed interest in elucidating the fundamental principles governing serine metabolism in all cells.

When exogenous serine is available, proliferating bacterial and mammalian cells deplete it rapidly, far faster than any other amino acid^9–11^. This observation has been cited as evidence that replicating cells require large amounts of serine to generate biomass and functional metabolites such as one-carbon units^2,12^. However, bacteria deaminate the majority of exogenous serine to produce pyruvate and ammonia^13,14^ (Fig 1A), even though this pathway competes with biomass-producing reactions and antagonizes growth^13^. Serine deamination is clearly important because *E. coli* expresses at least one of its three serine deaminase enzymes under any given condition^15^, yet the metabolic benefits(s) of this reaction remain unclear.

**Fig 1.**
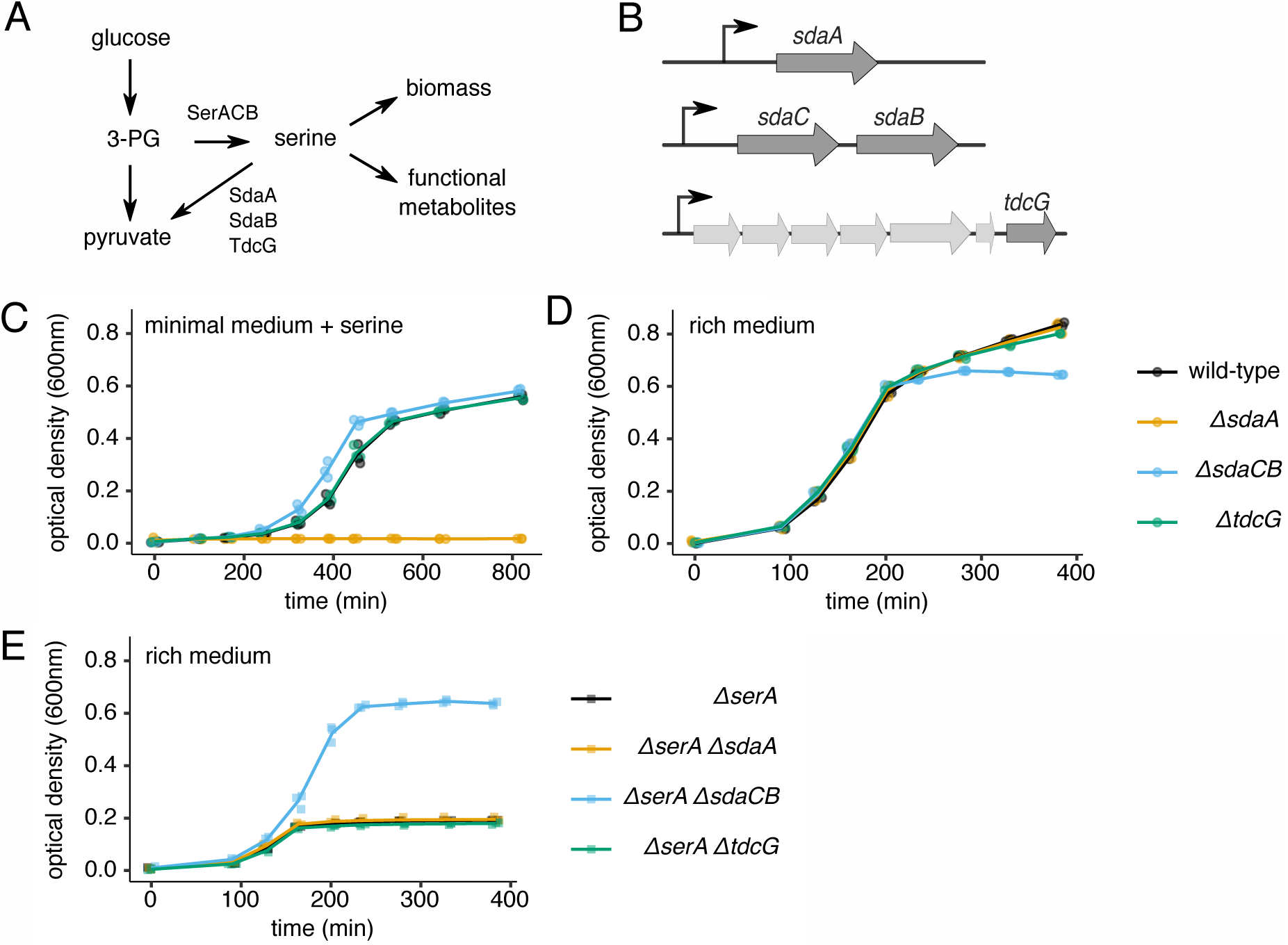
The role of serine deaminase operons in minimal vs. rich media. **(A)** Schematic of metabolic reactions and relevant enzymes involved in serine biosynthesis and utilization. 3-PG = 3-phosphoglycerate. **(B)** Schematic of the three loci in the *E. coli* genome that encode serine deaminase enzymes. **(C)** Growth curves of wild-type and serine deaminase single deletion strains grown in minimal medium containing 1.5mM serine. **(D)** Growth curves of wild-type and serine deaminase single deletion strains grown in rich medium containing 5mM serine. **(E)** Growth curves of *ΔserA* strains with or without a serine deaminase deletion grown in rich medium containing 5mM serine. For all growth curves, three biological replicates are shown as points with their averages connected by lines.

Serine deamination in bacteria is considered a detoxification mechanism because free intracellular serine inhibits biosynthesis of isoleucine and aromatic amino acids^16,17^. Indeed, all three *E. coli* serine deaminases have K_m_ values in the millimolar range^18,19^, consistent with a role in preventing serine excess^15^. *E. coli* cells unable to deaminate serine become severely elongated in the presence of exogenous serine, suggesting that high serine levels inhibit cell division^10,15^. This toxicity likely stems from defects in cell wall production given that mis-incorporation of serine in place of alanine into peptidoglycan crosslinks impairs peptidoglycan biosynthesis^20^. In all cases documented so far, toxicity is caused by import of excess serine; whether endogenously produced serine can become toxic is not known.

Bacteria do appear to actively avoid high intracellular serine levels. For example, the serine biosynthesis enzyme SerA is allosterically inhibited by serine^1,21^. Ribosome pausing is consistently observed at serine codons in bacterial ribosome profiling studies^22,23^, indicating that intracellular serine concentrations and/or charging levels of seryl-tRNAs are low compared to other amino acids. Surprisingly, the proportion of charged to uncharged seryl-tRNAs was observed to be much lower in rich media compared to minimal media (where exogenous serine is not available)^24^, suggesting that intracellular serine levels are lower in nutrient-rich conditions. Given that known serine toxicities should be less severe when all amino acids are provided, the physiological benefit(s) of maintaining low serine levels in rich media are unclear.

Here, we report that *E. coli* cells lacking the *sdaCB* operon, which encodes a serine transporter and a serine deaminase, lyse upon glucose depletion in rich media, thus revealing an unexpected role for *sdaCB* during adaptation to changes in nutrient availability. Importantly, lysis only occurs in the absence of exogenous serine, revealing for the first time that endogenously produced serine can be toxic. Heterologous expression of *sdaC* is sufficient to prevent lysis, implicating serine export, rather than deamination, in mediating the observed phenotype. Moreover, lysis can be modulated by altering glycine or alanine levels in the growth media, indicating the importance of maintaining a strict ratio of serine to other amino acids during metabolic shifts. Because glucose is the biosynthetic precursor of serine, we propose that SdaC is required to maintain serine homeostasis when cells are suddenly required to switch their serine production strategy.

## Results

### Cells lacking *sdaCB* exhibit a growth defect during entry into stationary phase

The *E. coli* genome encodes three separate serine deaminases (Fig 1B), of which at least one is always expressed; SdaA predominates in nutrient-poor conditions, SdaB predominates in nutrient-rich conditions, and TdcG predominates during anaerobiosis^15,25^. SdaB is co-expressed with the predicted serine transporter SdaC^26^, but the functional interplay between these two proteins is unknown. To investigate the role of serine deamination in different growth conditions, we constructed strains of *E. coli* (B**W**25113) harboring single deletions of *sdaA, sdaCB* or *tdcG* (Fig 1B). *E. coli* grown in minimal media containing serine require *sdaA* for survival^27,28^ because free serine inhibits the biosynthesis of some other amino acids^16,17^. We confirmed that our *ΔsdaA* strain is unable to grow in MOPS minimal medium containing 1.5mM serine (Fig 1C). Under the same conditions, deletion of the *sdaCB* or *tdcG* genes does not negatively impact growth (Fig 1C), consistent with reports that they are not highly expressed in minimal media^28^.

To investigate the role of serine deamination in nutrient-rich conditions, we monitored the growth of wild-type and the three serine deaminase single mutant strains in MOPS rich medium (minimal medium supplemented with amino acids and nucleobases) containing 5mM serine. All four strains exhibited identical growth kinetics until they began transitioning to stationary phase, at which point the optical density (OD) of the *ΔsdaCB* strain stopped increasing, resulting in a defect in overall growth yield (Fig 1D). This result suggested to us that *sdaCB* is not required for exponential growth in the presence of serine in rich media but may become beneficial when specific nutrient(s) are depleted. Interestingly, the growth yield defect of *ΔsdaCB* cells was smaller in a medium containing glycerol, rather than glucose, as the carbon source (Fig S1), suggesting that the effect may be glucose-specific.

To estimate the effect of SdaC/B activity on growth in rich media, we performed the same experiment described above in the context of a *ΔserA* background, in which cells are unable to synthesize serine *de novo.* The *ΔserA* strain depleted exogenous serine after three hours of growth, reaching a maximum growth yield of less than one third of wild-type cells (Fig 1E). While the growth yield of the *ΔserA ΔsdaA* and *ΔserA ΔtdcG* strains was the same as for the *ΔserA* strain, the *ΔserA ΔsdaCB* strain had a dramatically higher growth yield (Fig 1E), suggesting that SdaC/B activity competes with reactions that convert serine into biomass and consumes the majority of exogenous serine. Together, these results indicate that the *sdaCB* operon is highly active during nutrient-rich growth despite antagonizing biomass production, and may benefit cells as they transition to nutrient deprivation.

### Cells lacking *sdaCB* lyse upon glucose depletion

Because we observed a growth defect of the *ΔsdaCB* strain in glucose-containing rich medium during entry into stationary phase, we hypothesized that *sdaCB* might aid cells in making the metabolic changes necessary to adapt when glucose is exhausted. To test this idea, we compared the growth of wild-type and *ΔsdaCB* cells in modified rich media in which availability of glucose was limited. In media containing 1.4mM glucose (20-fold lower concentration than normal) and lacking serine, wild-type cells grew normally to a final OD_600_ that was approximately 2.5-fold lower than in rich medium (Fig 2A). In the same low glucose medium, the *ΔsdaCB* strain grew similar to the wild-type until approximately 150 min, at which point its optical density suddenly began decreasing, eventually dropping to background levels (Fig 2A). This phenotype was observed only when both glucose and serine were limiting; addition of either 5.6mM glucose or 3mM serine restored growth to wild-type levels (Fig 2A). Lysis of *ΔsdaCB* cells was also observed in a different *E. coli* genetic background (MG1655), indicating that the phenotype is not strain-specific (Fig S2).

**Fig 2.**
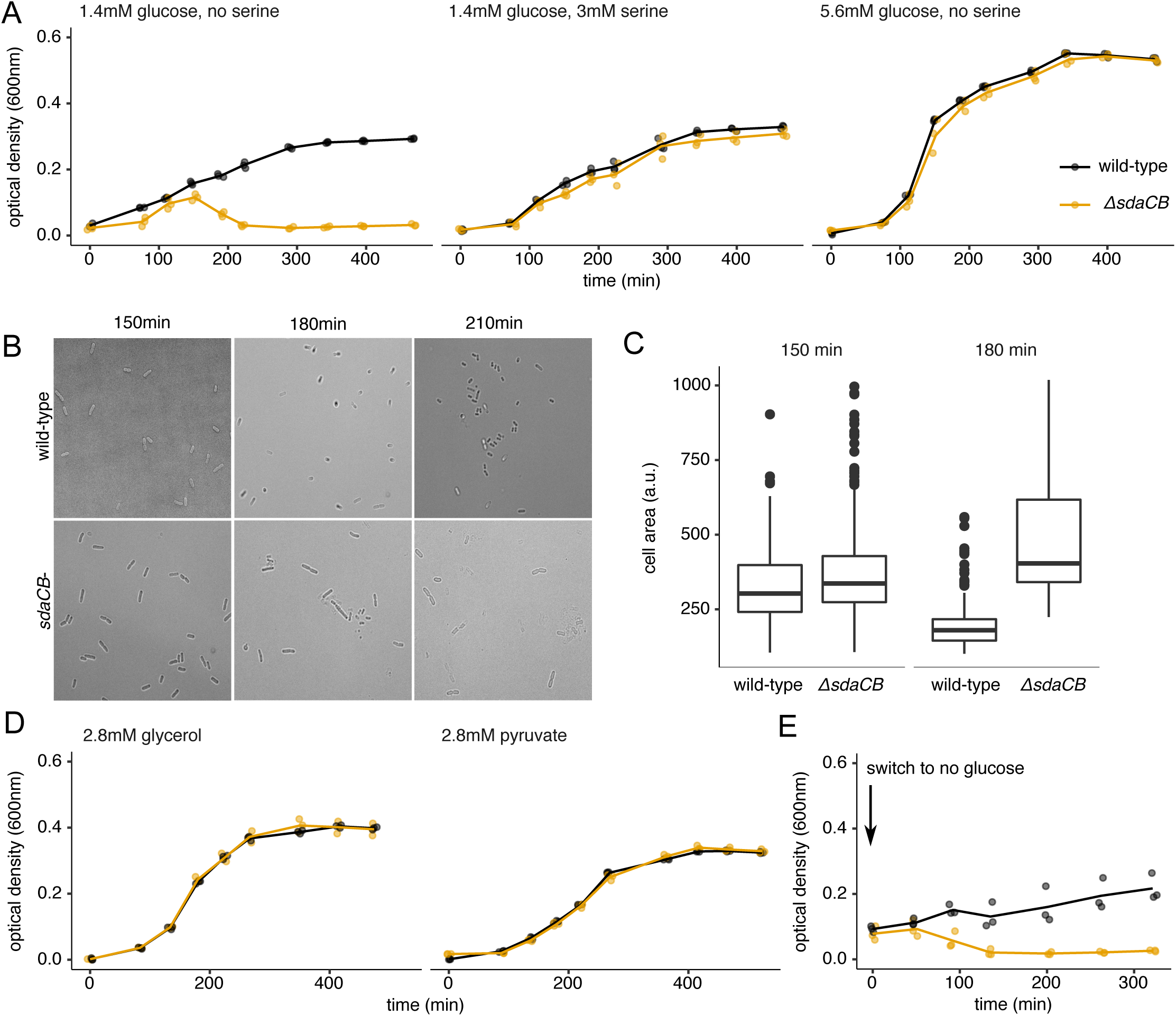
*sdaCB* prevents lysis upon glucose depletion. **(A)** Growth curves of wild-type and *ΔsdaCB* strains in modified rich medium containing 1.4mM (0.025%) glucose and no serine (left), 1.4mM glucose and 3mM serine (center), or 5.6mM glucose and no serine (right). **(B)** Bright-field microscopy images of wild-type and *ΔsdaCB* cells (60X magnification) grown as in (A) and transferred to poly-L-lysine-coated microscope slides at the indicated time points. **(C)** Individual cell areas measured as described in Methods for wild-type (n=206, 150 min; n=286, 180 min) and *ΔsdaCB* (n=316, 150 min; n=50, 180 min) strains grown as in (A). **(D)** Growth curves of wild-type and *ΔsdaCB* strains in modified rich medium (no glucose, no serine) containing 2.8mM (0.025%) glycerol or pyruvate. **(E)** Growth curves of wild-type and *ΔsdaCB* strains following removal of glucose. Overnight cultures were diluted 1:100 into modified rich medium (2.8mM glucose, no serine) for 105 min. The cultures were then spun down briefly to remove the media and cells were resuspended in modified rich medium (no glucose, no serine) prior to measurement of optical density over time. For all growth curves, three biological replicates are shown as points with their averages connected by lines.

A downward trend in optical density is often indicative of cell lysis; indeed, we observed by microscopy that *ΔsdaCB* cells, but not wild-type cells, lysed approximately 150 min after dilution of the overnight cultures (Fig 2B). Interestingly, lysis was not preceded by significant morphological changes (Fig 2B), as has been observed in other studies of serine deaminase mutants^10,15^, but we did observe a decrease in cell size – as expected for entry into stationary phase^29^ – only for the wild-type strain (Fig 2C). The lysis phenotype did not occur when glucose was replaced by alternative carbon sources, such as glycerol or pyruvate (Fig 2D), suggesting a specific role for glucose depletion in causing lysis of *ΔsdaCB* cells. To test directly whether glucose depletion is the trigger for lysis of the *ΔsdaCB* strain, we grew cells in MOPS rich medium containing 2.8mM glucose and no serine for 100 min and then switched them to the same medium containing no glucose. As predicted, the medium switch caused complete lysis of the *ΔsdaCB* strain, but not the wild-type strain (Fig 2E). This result supports the conclusion that *ΔsdaCB* cells lyse specifically upon glucose depletion in the absence of exogenous serine.

### *sdaC* is sufficient to prevent lysis

To gain insight into the mechanism by which loss of *sdaCB* causes lysis, we measured the individual effects of *sdaC* and *sdaB* deletions on the lysis phenotype in medium with low glucose and no serine. Unexpectedly, the serine deaminase mutant, *ΔsdaB,* did not lyse upon glucose depletion, whereas the serine transporter mutant, *ΔsdaC,* did lyse (Fig 3A). qRT-PCR results revealed that deletion of *sdaC* modestly reduced *sdaB* expression(Fig S3A), likely because *sdaC* is upstream of *sdaB* in the operon. To confirm that lysis of *ΔsdaC* cells was caused by loss of *sdaC* alone, rather than disrupted expression of both *sdaC* and *sdaB,* we tested whether low-copy plasmids^30^ (pSC*101 *ori,* 3–4 copies/cell) expressing *sdaC* or *sdaB* alone could prevent lysis. A plasmid expressing *sdaC* under the control of its native promoter completely rescued lysis of the *ΔsdaCB* strain (Fig 3B), whereas *ΔsdaCB* cells still lysed when harboring a plasmid expressing *sdaB* under the control of the *sdaCB* promoter (Fig 3B). Together, these results indicate that *sdaC* expression is sufficient to prevent lysis, regardless of whether *sdaB* is present (Fig 3B).

**Fig 3.**
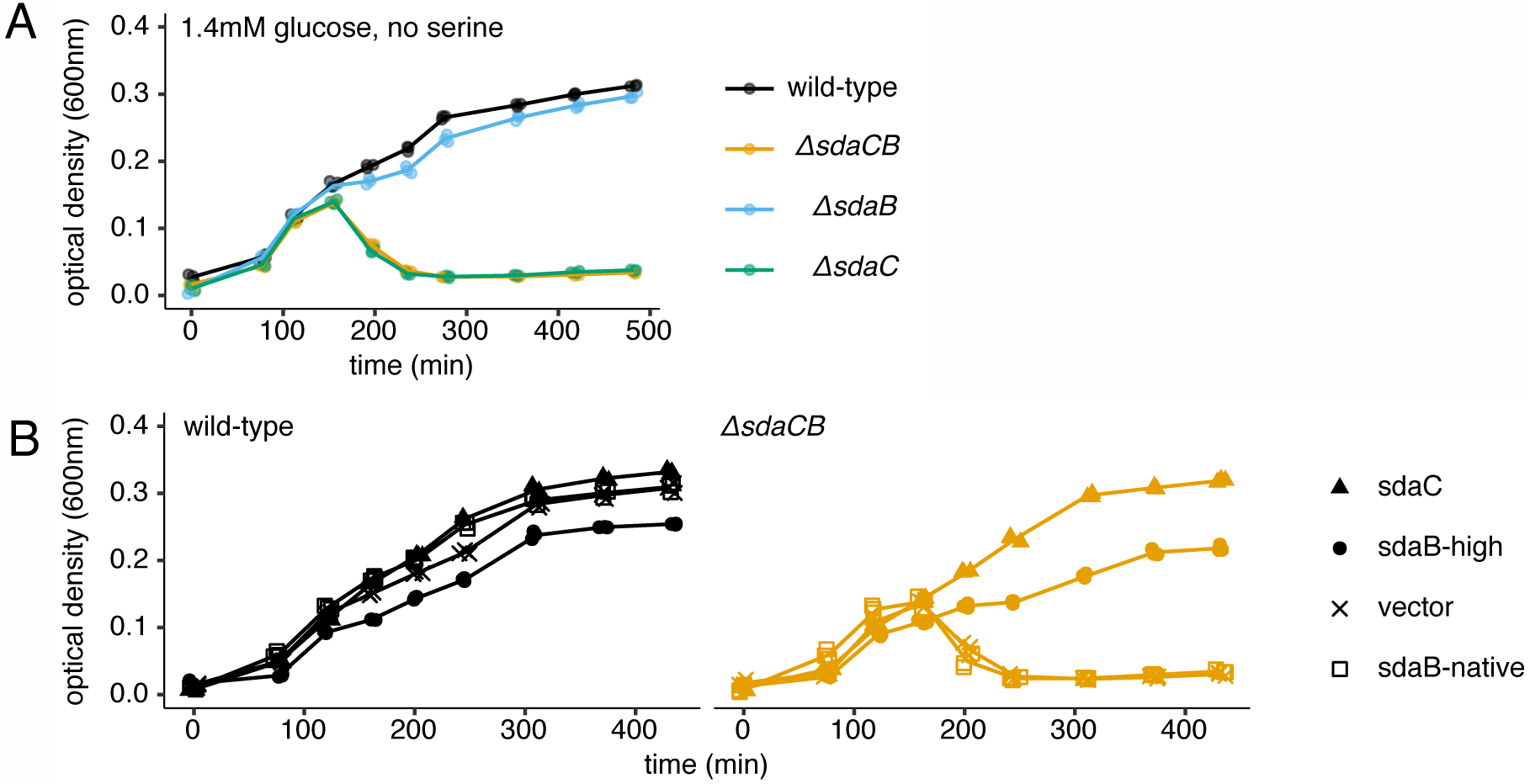
*sdaC* is sufficient to prevent lysis. **(A)** Growth curves of wild-type, *ΔsdaC, ΔsdaB* and *ΔsdaCB* strains in modified rich medium (1.4mM glucose, no serine). **(B)** Growth curves of wild-type and *ΔsdaCB* strains harboring a low-copy plasmid expressing *yfp* (vector) or derivatives expressing *sdaC* under the control of its native promoter and leader sequence (sdaC), *sdaB* under the control of its native promoter (sdaB-native) or *sdaB* under the control of the strong promoter pLtetO-1 (sdaB). Cells were grown in the same medium as in A. For all growth curves, three biological replicates are shown as points with their averages connected by lines.

Interestingly, a plasmid expressing *sdaB* under the control of the highly active pLtetO-1 promoter^30^ did prevent lysis of the *ΔsdaCB* strain (Fig 3B), indicating that over-expression of *sdaB* can substitute for loss of *sdaC*. To rule out the possibility that insufficiency of the *ΔsdaB* mutation to cause lysis is due to compensatory induction of another serine deaminase, we constructed a *ΔsdaB ΔsdaA ΔtdcG* triple deletion strain; this strain also did not lyse upon glucose depletion (Fig S3B). Finally, to rule out the possibility that a genetic determinant other than the *sdaC* coding region is responsible for preventing lysis, we constructed a derivative of the *sdaC*-expressing plasmid containing a point mutation that introduces a premature stop codon after the 82nd amino acid of *sdaC;* this plasmid was unable to prevent lysis of *ΔsdaCB* cells (Fig S3C).

SdaC is thought to be a proton-serine symporter that imports serine under physiological conditions^26,31^. However, our observations that (i) *sdaC* is required for survival in a condition in which there is no exogenous serine and (ii) over-expression of a serine-consuming protein (SdaB) can fulfill the same biological function as SdaC led us to consider that serine export may be the relevant function of SdaC under our experimental conditions. To directly test whether SdaC exports serine under physiological conditions, we grew wild-type and *ΔsdaC* cells to mid-exponential phase in rich medium containing no serine and performed mass spectrometry on culture supernatants to quantify metabolites. If SdaC exports serine under this condition, we expected to observe serine in wild-type supernatants but not in *ΔsdaC* supernatants or sterile medium. However, slightly more serine was detected in the *ΔsdaC* supernatants (~25µM) than in wild-type supernatants (~10µM) or sterile medium (~6µM) (Fig S4). None of the other amino acids was significantly different between wild-type and mutant supernatants (Fig S4). These results are not consistent with serine export by SdaC during glucose-replete growth, but it remains possible that glucose depletion in the lysis condition reverses the electrochemical gradient for serine transport through SdaC given that glucose increases the proton motive force^32–34^ and SdaC is a proton/serine symporter. We were unable to test this possibility directly by measuring serine levels in the medium during glucose depletion since intracellular serine released from lysed cells would confound our measurements.

### The glycine cleavage system promotes lysis upon glucose depletion

*De novo* serine biosynthesis begins with the glycolytic intermediate 3-phosphoglycerate, which is converted to serine via three steps^1,6,21^ (Fig 1A). Because lysis of the *ΔsdaC* strain occurs when cells cannot obtain serine from the growth medium and have run out of its biosynthetic precursor (glucose), we reasoned that lysis may coincide with induction of alternative pathways for serine production

(Fig 4A). The primary means of generating serine besides *de novo* synthesis is to catabolize other amino acids, such as glycine or threonine^1^; thus, it could be that amino acid catabolism contributes to the lysis phenotype. In accordance with this hypothesis, the *ΔsdaC* strain did not lyse when glycine was omitted from the lysis-promoting growth medium (1.4mM glucose, no serine; Fig 4B).

**Fig 4.**
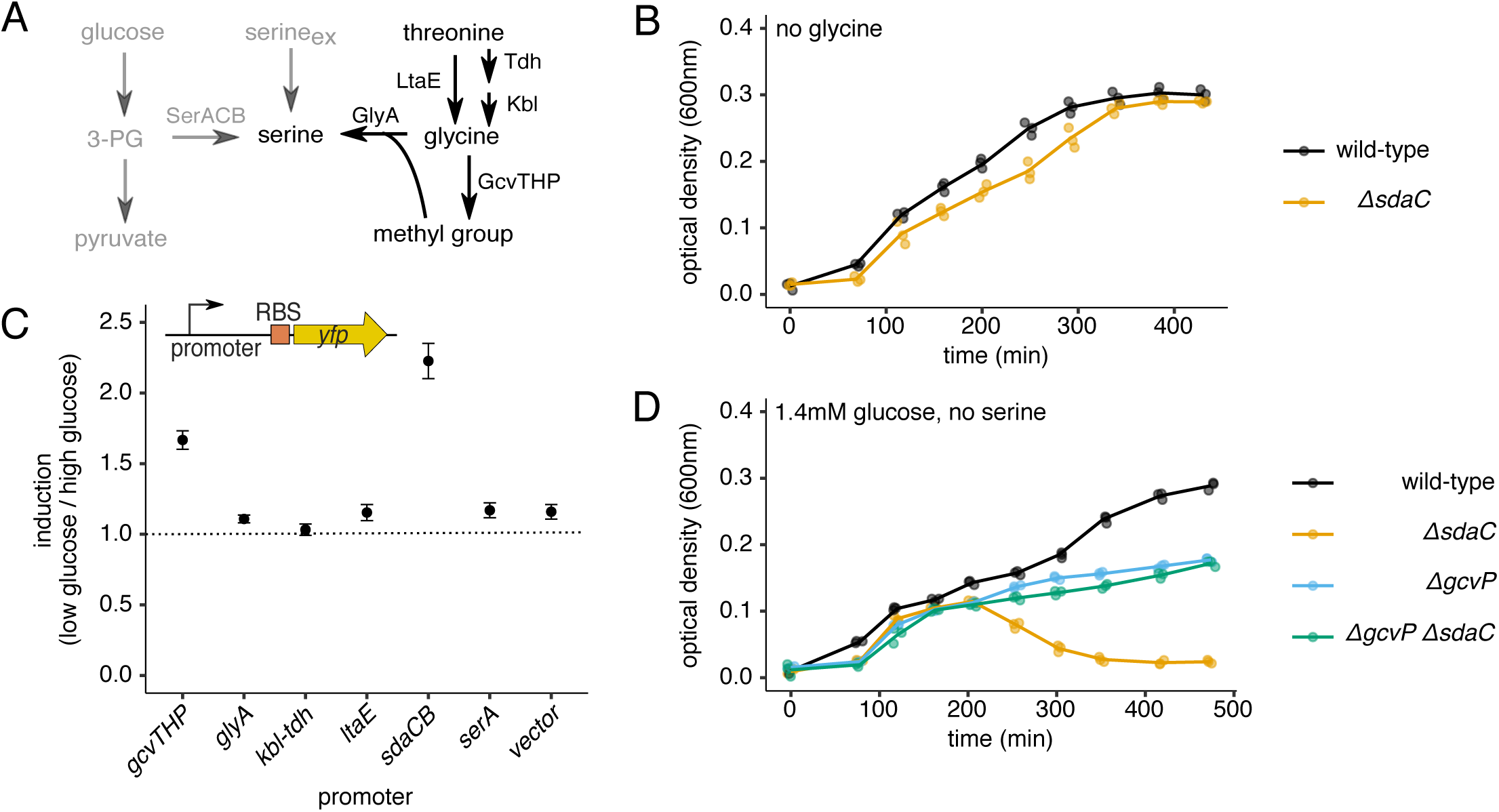
The glycine cleavage system promotes lysis upon glucose depletion. **(A)** Schematic of metabolic pathways involved in serine production. **(B)** Growth curves of wild-type and *ΔsdaC* strains in modified rich medium (1.4mM glucose, no serine, no glycine). **(C)** Relative YFP fluorescence (glucose-depleted vs. glucose-containing media) from wild-type strains harboring transcriptional reporters for genes related to glycine and threonine catabolism. Raw fluorescence values were first normalized to culture OD, and relative fluorescence was calculated by dividing the YFP/OD signal (averaged over the three time points directly after glucose depletion occurs in the 1.4mM glucose condition) from cells grown in media containing 1.4mM vs. 5.6mM glucose. **(D)** Growth curves of wild-type, *ΔgcvP, ΔsdaC* and *ΔgcvP ΔsdaC* strains in modified rich medium (1.4mM glucose, no serine). For all growth curves, three biological replicates are shown as points with their averages connected by lines. For YFP induction, average and standard error over biological triplicates is shown.

In order to determine which biochemical pathway(s) involved in serine production might become more active upon glucose depletion, we constructed transcriptional reporters for operons known to be involved in generating serine from other amino acids: the glycine cleavage system *(gcvTHP),* serine hydroxymethyltransferase (*glyA*), threonine dehydrogenase *(kbl-tdh)* and threonine aldolase (*ltaE*) (Fig 4A). As controls, we also constructed transcriptional reporters for the serine deaminase operon *sdaCB* and the serine biosynthesis gene *serA.* Promoters of interest were cloned upstream of the *yfp* translation initiation signals (RBS) and coding region in a low copy plasmid (Fig 4C). We measured YFP levels over time in wild-type strains harboring each reporter in two conditions: the lysis promoting medium (1.4mM glucose, no serine) or the same medium containing additional glucose (5.6mM). We compared YFP production from a given reporter over the time range at which *ΔsdaC* cells lyse in the low-glucose condition (Fig 4C); cells grown in the higher glucose condition have not yet depleted the glucose supply by this time. The only amino acid catabolic operon that showed transcriptional induction upon glucose depletion was *gcvTHP* (Fig 4C), which encodes enzymes that extract a methyl group from glycine. This methyl group can then be combined with an additional glycine molecule by GlyA to produce serine^1^. Of note, we also observed induction of *sdaCB* itself, consistent with the known activation of this operon by cyclic AMP^35^ (Fig 4C).

Based on the role of glycine in promoting lysis (Fig 2A vs. Fig 4B) and the transcriptional induction of the glycine cleavage operon upon glucose depletion (Fig 4C), we hypothesized that glycine cleavage by GcvTHP triggers lysis of the *ΔsdaC* strain. To test this idea, we deleted the *gcvP* gene, encoding glycine decarboxylase, in both wild-type and *ΔsdaC* backgrounds. In accordance with our hypothesis, *gcvP* deletion prevented lysis of the *ΔsdaC* strain (Fig 4D). Notably, *ΔgcvP* strains exhibited a growth defect relative to wild-type following glucose depletion (Fig 4D), strengthening the notion that glycine cleavage is a major means of satisfying metabolic requirements when serine and glucose are absent.

### The lysis phenotype is associated with a high serine to alanine ratio

We next sought to understand why conversion of glycine to serine upon glucose depletion promotes cell lysis. In bacteria, lysis is often indicative of a defect in cell wall biogenesis. A major component of the bacterial cell wall is peptidoglycan, which is composed of glycan chains crosslinked by short peptides. In *E. coli* and many other bacteria, the peptide crosslinks have a high alanine content^36^. It was recently shown that serine can be mis-incorporated into peptidoglycan crosslinks in place of alanine^20^, likely due to weak substrate specificity of the enzyme MurC, which attaches L-alanine to UDP-N-acetylmuramic acid in an early step of peptidoglycan biosynthesis^37^. Mis-incorporation of serine was shown to reduce the rate of peptidoglycan production and sensitize cells to *β*-lactam antibiotics^20^. As a result, we hypothesized that lysis of the *ΔsdaC* strain could be due to a higher than normal serine to alanine ratio (Fig 5A). Indeed, when we added 1.8mM alanine to the lysis-promoting growth medium (instead of the standard 0.8mM in our MOPS rich medium), *ΔsdaC* cells no longer lysed (Fig 5B). Strikingly, omission of alanine from the lysis-promoting medium led to lysis of the wild-type strain (Fig 5C), further implicating a high serine to alanine ratio in triggering lysis upon glucose depletion. We also tested whether an orthogonal approach to altering alanine availability promotes lysis; indeed, deletion of one of *E. coli’s* three alanine-synthesizing transaminases, *alaA,* resulted in lysis of cells in which *sdaCB* was intact (Fig 5D). Finally, addition of equimolar pyruvate to the lysis-promoting medium was insufficient to prevent lysis of *ΔsdaC* cells, reinforcing the notion that serine removal, rather than a product of serine deamination, is required in this condition (Fig 5E).

**Fig 5.**
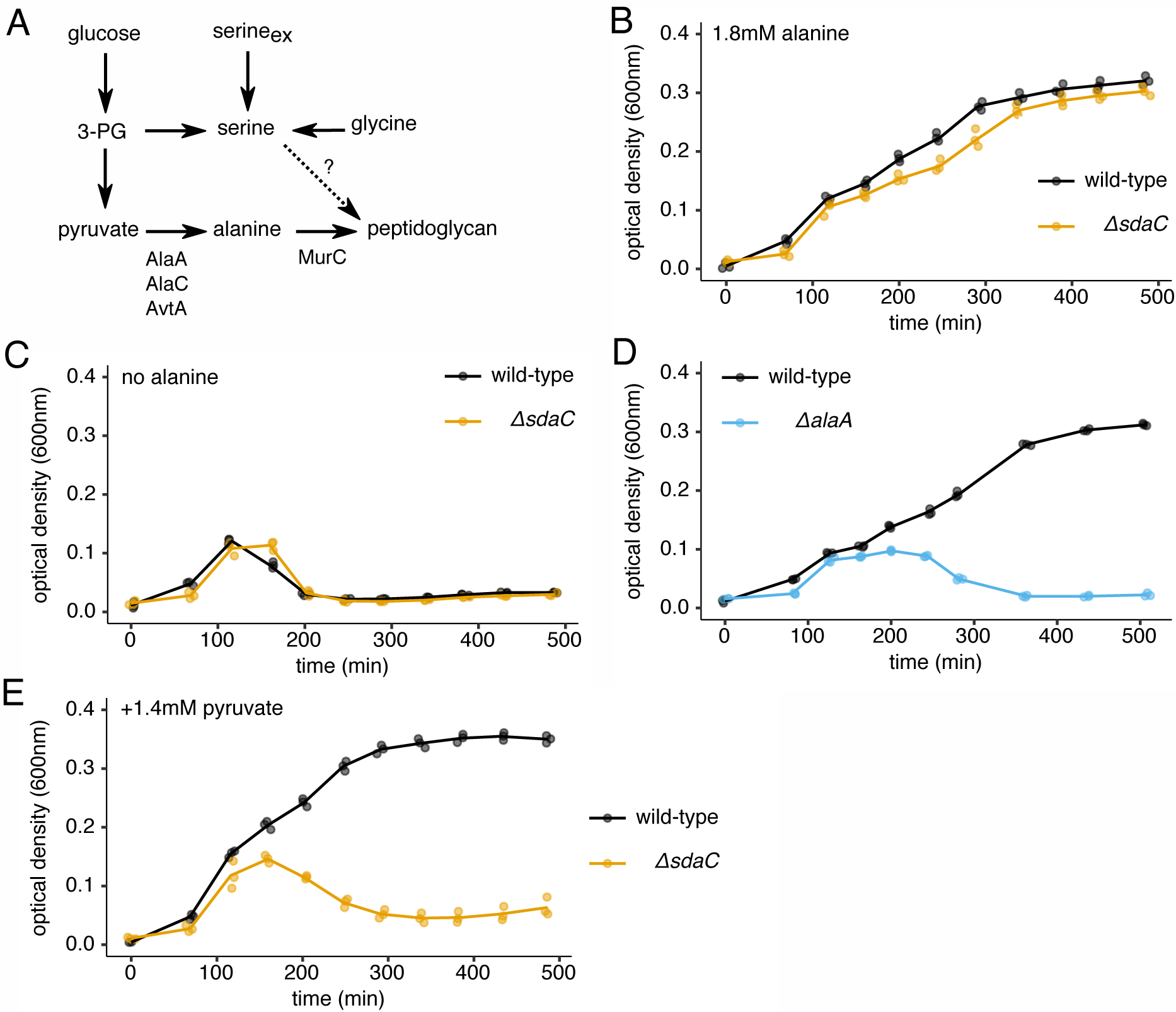
The lysis phenotype is associated with a high serine to alanine ratio. **(A)** Schematic of selected reactions related to alanine and serine metabolism. **(B)** Growth curves of wild-type and *ΔsdaC* strains in modified rich medium (1.4mM glucose, no serine, 1.8mM alanine). **(C)** Growth curves of wild-type and *ΔsdaC* strains in modified rich medium (1.4mM glucose, no serine, no alanine). **(D)** Growth curves of wild-type and *ΔalaA* strains in modified rich medium (1.4mM glucose, no serine, 0.8mM alanine). **(E)** Growth curves of wild-type and *ΔsdaC* strains in modified rich medium (1.4mM glucose, no serine, 0.8mM alanine, 1.4mM pyruvate). For all growth curves, three biological replicates are shown as points with their averages connected by lines.

## Discussion

In this work, we found that glucose depletion in the absence of serine causes lysis of *E. coli* cells lacking the *sdaCB* operon. Our results suggest that glucose depletion triggers catabolism of glycine, and possibly other amino acids, to satisfy cellular serine requirements and that the resulting increase in intracellular serine levels is surprisingly deleterious (Fig 6). In this context, serine detoxification is performed by the gene products of the *sdaCB* operon, as evidenced by induction of *sdaCB* expression (Fig 4C) and lysis of *ΔsdaCB* cells (Fig 2A) upon glucose depletion. Interestingly, we found SdaC to be the dominant activity in preventing toxicity of intracellularly produced serine, thus revealing an unexpected role for serine transport in adaptation to changes in nutrient availability. To our knowledge, this is the first report that intracellularly produced serine can be toxic and that a serine transporter can mitigate serine toxicity. The lysis phenotype is dependent on a high serine to alanine ratio, suggesting that serine toxicity in this context arises due to interference of free serine with the alanine-dependent step of peptidoglycan biogenesis, as reported by other recent studies^10,15,20^.

**Fig 6.**
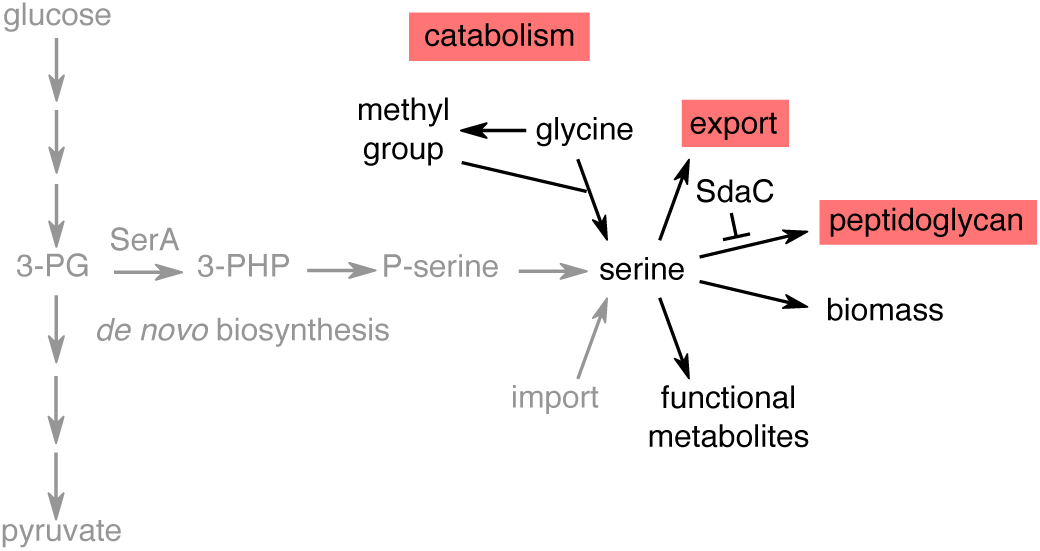
Model for serine metabolism and toxicity upon glucose depletion. During exponential growth, intracellular serine synthesized from glucose or imported from the medium is efficiently funneled to reactions that produce biomass and/or functional metabolites. Upon glucose depletion in the absence of exogenous serine, cells must shift to amino acid catabolism as a serine source. This abrupt switch causes a transient spike in free serine levels that can lead to toxic incorporation of serine into peptidoglycan if the cell is unable to export excess serine via SdaC.

Our observations that the presence of glycine and activity of the glycine cleavage system promote cell lysis upon glucose depletion (Fig 4) are consistent with a model in which a switch to glycine catabolism produces a sudden and potentially toxic increase in serine levels. Although the direct product of the glycine cleavage system is a methyl group rather than serine, this methyl group is used by GlyA to convert additional glycine molecules into serine. We were unable to perform experiments in a *ΔglyA* background due to the extremely slow growth rate of strains harboring this deletion. We conclude that toxicity is due to excess serine, rather than a lack of serine deamination products, because (i) a mutant lacking all three serine deaminases did not lyse (Fig S3B) and (ii) exogenous pyruvate was unable to prevent lysis of *ΔsdaC* cells (Fig 5E).

Interestingly, *ΔserA* cells grown in glucose-replete media do not appear to activate amino acid catabolic pathways upon exhaustion of exogenous serine, even though this should allow them to continue growing (Fig 1E). This situation, which is presumably due to glucose-mediated repression^38^, makes sense from an evolutionary perspective because wild-type cells can produce serine from glucose, and amino acid precursors for producing serine are themselves useful as substrates for protein synthesis. However, our observation also implies that cells strongly prefer *de novo* synthesis to amino acid catabolism for endogenous production of serine. A possible explanation for this preference is that *de novo* synthesis provides a more controlled environment, for instance due to tight control of reaction rates, with less potential for toxicity of free serine compared to the *gcvTHP/glyA* pathway.

We were surprised to find that *sdaC* is sufficient to prevent the lysis phenotype, even in a strain lacking all three serine deaminases (Fig S3B). SdaC is a proton/serine symporter that has been shown to import serine under physiological conditions^26,31^, yet the medium that triggers lysis of *ΔsdaC* cells is devoid of exogenous serine. Thus, we propose that serine export is the function of SdaC that becomes essential upon glucose depletion. This notion is bolstered by our observation that strong over-expression of the serine deaminase SdaB, which effectively removes serine from the cell, can compensate for loss of SdaC. The other known serine transporters in *E. coli* are unlikely to export serine in the lysis-promoting condition; the sodium-serine symporter SstT would require a reversal of the sodium gradient and the anaerobic proton-threonine/serine transporter TdcC is not expressed^39^.

Our metabolomic analysis of wild-type and *ΔsdaC* culture supernatants did not provide direct evidence of serine export by SdaC, but this experiment was performed in glucose-replete conditions. Because SdaC is a proton/serine symporter and intracellular pH is likely to drop upon glucose depletion^32–34^, we suspect that glucose depletion shifts the electrochemical gradient towards proton/serine co-export. An alternative possibility is that SdaC is required to scavenge serine released from dead cells and/or exported by another protein, but our results from modulating glycine and alanine availability are not consistent with serine starvation being the trigger for cell lysis. SdaC could also possess an additional serine detoxification function independent of its serine transport activity, but we consider this possibility unlikely due to the high homology of SdaC to well-defined amino acid transporters such as TdcC^26^.

Recent studies of *E. coli* lacking all three serine deaminases found that a triple mutant strain forms large, misshapen cells in glucose-replete minimal media containing exogenous serine^10,15^. These serine-induced morphology defects occured in the presence of *sdaC,* whereas we observe lysis of *ΔsdaC* cells but not *ΔsdaB ΔsdaA ΔtdcG* cells in our experimental condition (Fig S3B). This difference may be due to the source and amount of excess serine being different; in the case documented by Zhang et al, serine deaminases detoxify exogenous serine (presumably the presence of glucose and serine in the medium makes the electrochemical gradient less favorable for serine export by SdaC), whereas in our case, endogenously produced serine (whose concentration is likely lower than the millimolar K_m_ of the serine deaminases) can be detoxified by serine export through SdaC (which is more favorable due to both a lack of external serine and a low internal pH caused by glucose depletion).

Consistent with our observation that modulating alanine availability influences the lysis phenotype (Fig 5), Zhang et al reported that serine toxicity could be prevented by alanine supplementation or over-expression of the alanine ligase, MurC^10^. Their hypothesis that free serine inhibits cell wall biogenesis by competing with alanine for MurC^10^ was recently corroborated by HPLC analysis directly demonstrating incorporation of serine into peptidoglycan crosslinks at the position normally occupied by alanine when intracellular serine levels are high^20^. Serine incorporation was associated with a reduced rate of peptidoglycan production and sensitivity to *β*-lactam antibiotics^20^, indicating that the presence of serine in peptidoglycan crosslinks is indeed deleterious. Thus, we propose that in our experimental condition, glucose depletion causes a sudden imbalance between serine and alanine levels that can lead to toxic incorporation of serine into the cell wall.

More broadly, our findings can inform efforts to improve genome-scale models of metabolism, which typically use biomass production as the sole objective function and thus do not account for potential toxicity of metabolites present in excess^40^. Given the importance of serine detoxification for survival of bacteria under a variety of conditions, future iterations of metabolic models may need to incorporate this principle, possibly by requiring serine export and/or deamination. In addition, our results suggest that targeting SdaC might be an effective strategy for increasing the efficacy of *β*-lactam antibiotics.

## Acknowledgments

We thank Heungwon Park and Julio Vasquez for assistance with microscopy experiments, and Rob Pepin and Dan Raftery at the Northwest Metabolomics Research Center for conducting the metabolomic analysis of culture supernatants. This work was supported by grant R35 GM119835 of the NIGMS (NIH) to ARS. MAK was supported by the Fred Hutch Chromosome Metabolism and Cancer training grant, NIH-2T32CA9657-26A1.

## Author Contributions

MAK: Conceptualization, Investigation, Formal Analysis, Writing – Original Draft Preparation; ARS: Conceptualization, Writing – Review & Editing, Supervision

## Materials and Methods

### Data and code availability

Raw data and scripts for performing all analyses and generating figures in this manuscript are publicly available at https://github.com/rasilab/mkriner_2018.

### Growth media

Bacteria were grown in a MOPS-based rich defined medium^41^ (adapted from the *E. coli* genome project^42^), containing 0.5% (28mM) glucose as the carbon source unless otherwise noted. The 5X Supplement was replaced with a 5X solution containing 4mM of each amino acid (0.8mM final concentration) except serine, glycine and alanine, which were added separately. When needed, glycine and alanine were added to a final concentration of 0.8mM and serine was added to a final concentration of 10mM, unless otherwise noted. The medium that produces lysis of *ΔsdaC* strains contains 0.025% (1.4mM) glucose and no serine. For all growth curve experiments in rich media, overnight cultures were grown in rich medium containing 0.5% glucose and 10mM serine and diluted 1:100 into experimental media. For the minimal medium experiment (Fig 1C), nucleobases and amino acids were omitted. Overnight cultures were grown in minimal medium without serine and diluted 1:100 into minimal medium containing 1.5mM serine. For cloning and strain construction, bacteria were grown in LB medium (BD/Difco) and the antibiotics carbenicillin (Carb; 100 µg/ml), chloramphenicol (Cm; 25 µg/ml) or kanamycin (Kan; 25 µg/ml) were included when required.

### Strain construction

All experiments were carried out with wild-type *Escherichia coli* strains BW25113^43^ or MG1655^44^ and mutant derivatives constructed using the one-step gene disruption method^43,45^ (Table S1). Briefly, a Kan or Cm antibiotic cassette was amplified from plasmid pKD13 or pKD32, respectively, using primers containing gene-specific 5**’** extensions (Table S2). Each cassette was designed to remove the desired coding region(s) upon homologous recombination, leaving regulatory regions intact. Recombination was achieved by electroporating 500ng of cassette DNA into a strain harboring plasmid pSIM6. pSIM6-harboring competent cells were prepared following growth to OD_600_=0.4−0.5 in LB and heat shock at 42°C for 15 min to induce expression of the λ-Red recombinase machinery^45^. When desired, the antibiotic resistance cassette was excised using pCP20 as described^43^, leaving an 84bp scar sequence. When making additional mutations in a strain already harboring a cassette and/or scar sequence, the antibiotic cassette was generated by colony PCR from a strain already containing the insertion using primers approximately 100 bp outside of the insertion. This strategy yields a cassette with longer homology arms and significantly reduced integration of the cassette into the locus of the first mutation. Successful strain construction was confirmed by colony PCR and Sanger sequencing.

### Plasmid construction

pASEC1 (Addgene plasmid #53241), a very low copy expression vector (SC*101 ori, 3–4 copies per cell) used in our previous work^46^, was used as a backbone for constructing the transcriptional reporters shown in Fig 4C. For each gene of interest, the promoter region was amplified from genomic DNA using the primers listed in Table S2 and inserted between the XhoI and EcoRI sites of pASEC1 via Gibson assembly. This strategy removes the pLtetO-1 promoter and places the gene-specific promoter directly upstream of the T7 ribosome binding site and *yfp* coding region. Successful construction was confirmed by colony PCR and Sanger sequencing.

### Growth and fluorescence measurements

Bacterial strains were streaked onto LB plates and single colonies were used to inoculate overnight cultures of MOPS rich medium (with 10mM serine) in triplicate in standard 96-well plates (Costar 3595). Cultures were incubated overnight at 37°C with shaking (Titramax 100 shaker, 900 rpm), diluted 1:100 into experimental media, and returned to 37°C with shaking. Cell density (absorbance at 600 nm) and/or YFP fluorescence (excitation 504 nm and emission 540 nm) were monitored every 40-60 min using a Tecan plate reader (Infinite M1000 PRO). To minimize evaporation, wells on the edge of the plate were filled with water and plates were sealed with parafilm between readings.

### Microscopy

Cells harboring pASEC1 were grown in 96-well plates as described above. At the indicated time points, 3µl of culture was transferred to a poly-L-lysine-coated microscope slide and covered with a coverslip. Cells were imaged on a DeltaVision Elite microscope at 60X magnification in the bright field and YFP channels. Cell number and cell area were measured using ImageJ software. YFP channel images (which only capture live cells) were converted to 16-bit grayscale and the threshold was adjusted to 61-255 to remove background. Cell area was calculated using the Analyze Particles function. Six fields per strain were imaged and quantified at each time point.

### Quantitative PCR

3ml cultures were grown to mid-exponential phase in MOPS rich medium (with 5mM serine) and harvested by centrifugation at 3000 g for 5 min. Cell pellets were resuspended in 250µl 0.3M sodium acetate and 10 mM EDTA (pH 4.5) and mixed with 250µl acid phenol-chloroform (pH 4.5) and 250µl glass beads (G1877, Sigma). Samples were vortexed for 3 min at maximum speed and clarified by centrifugation at 12,000 g for 10 min at 4°C. The aqueous layer was collected and RNA was purified using the Direct-zol RNA purification kit (Zymo Research).

300 ng purified RNA was treated with DNaseI (NEB) for 10 min at 37°C according to the manufacturer’s instructions. Reverse transcription (RT) was performed using Maxima RT enzyme (EP0741, Thermo Fisher) and random hexamer primers using a 20µl reaction volume. After incubation at 25°C for 10 min, 50°C for 30 min and 85°C for 5 min, samples were diluted 10-fold and 2 µl of diluted sample was used as template for a 10 µl qPCR reaction in the next step. qPCR was performed using Maxima SYBR Green/ROX qPCR Master Mix (FERK0221, Thermo Fisher) according to the manufacturer’s instructions. qPCR was performed in duplicate for each RT reaction using primers listed in Table S2. Negative RT controls were included to confirm the absence of DNA contamination. For each gene, _CT_ values were converted to mRNA quantity by generating a standard curve from genomic DNA. *gapA* mRNA was used as internal reference to normalize all other mRNA levels.

### Metabolomics

Wild-type and *ΔsdaC* overnight cultures grown in MOPS rich medium (with 10mM serine) were diluted 1:100 into MOPS rich medium (without serine) and grown at 37°C with shaking. Upon reaching early exponential phase (OD_600_=0.3), supernatants were isolated by passing cultures through a 0.22µm filter. Targeted LC-MS/MS was performed on a system consisting of Shimadzu Nexera XR LC-20AD pumps coupled to a Sciex 6500+ triple quadrupole spectrometer operating in scheduled MRM detection mode through the Sciex Analyst 1.6.3 software. The samples were separated on a Waters Xbridge BEH amide column operated in a HILIC regime. Metabolite concentrations were quantified using Multiquant 3.0 software. Absolute quantitation of all amino acids was possible through comparison to stable isotope-labeled internal standards.

## Supplementary Figures

**Fig S1.**
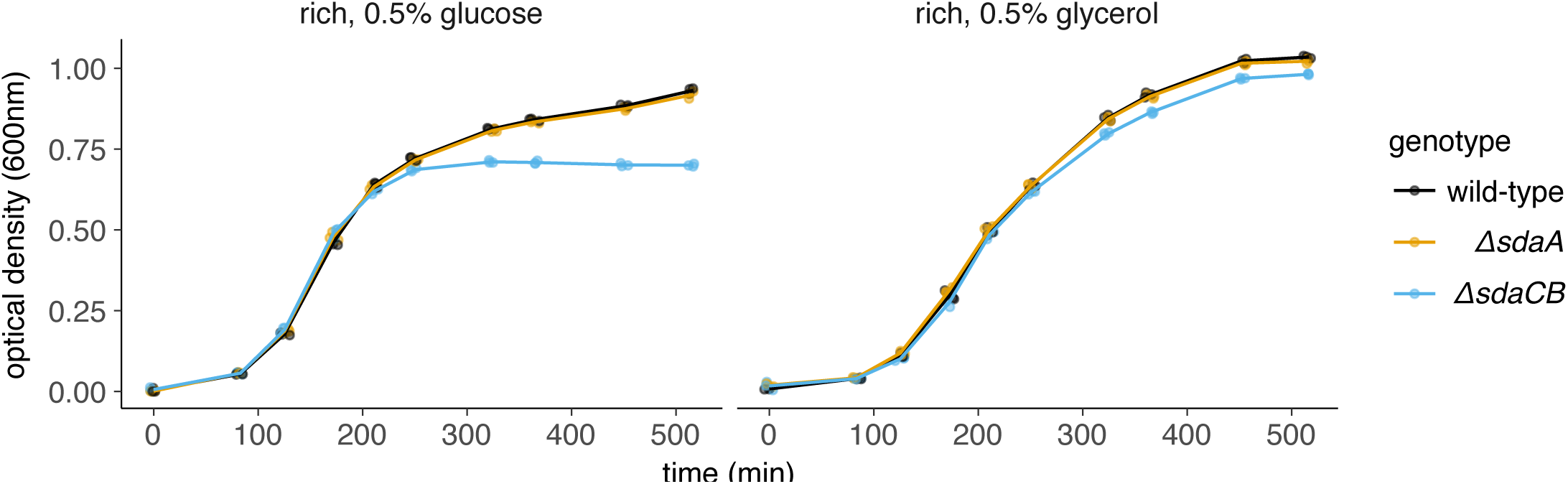
*ΔsdaCB* growth defect is specific to glucose-containing media. Growth curves of wild-type and serine deaminase single deletion strains grown in rich medium containing 5mM serine and 0.5% (56mM) glycerol or 0.5% (28mM) glucose as the carbon source. Three biological replicates are shown as points with their averages connected by lines.

**Fig S2.**
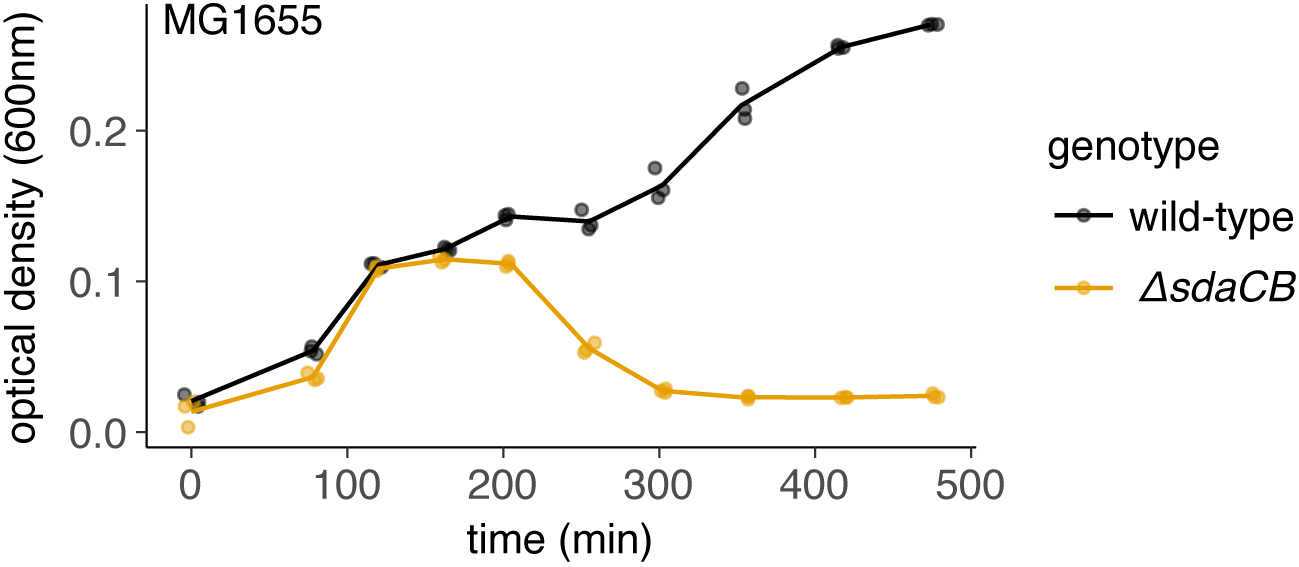
The lysis phenotype also occurs in the MG1655 genetic background. Growth curves of MG1655 and MG1655 *ΔsdaCB* strains in rich medium containing 0.025% (1.4mM) glucose and no serine. Three biological replicates are shown as points with their averages connected by lines.

**Fig S3.**
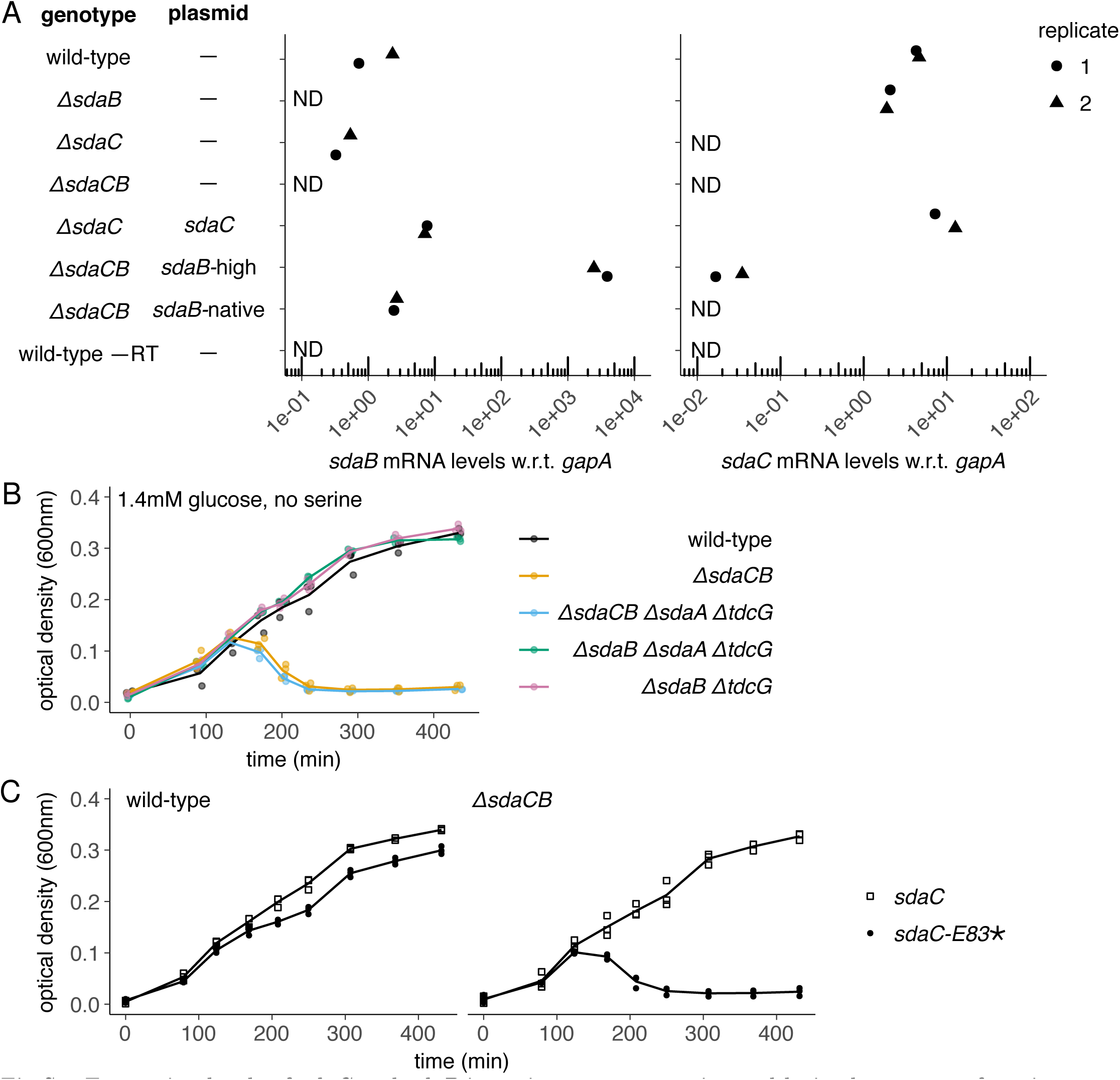
Expression levels of *sdaC* and *sdaB* in various mutant strains and lysis phenotypes of strains harboring different combinations of serine deaminase operon deletions and/or plasmids. **(A)** mRNA levels, as measured using qRT-PCR, of indicated strains grown to mid-exponential phase in MOPS rich medium containing 5mM serine. mRNA levels for *sdaC* and *sdaB* were normalized to the control gene *gapA.* ND = not detected, −RT = no reverse transcriptase. **(B)** Growth curves of strains harboring mutations in 0-3 of the serine deaminase loci in rich medium containing 0.025% (1.4mM) glucose and no serine. **(C)** Growth curves of wild-type and *ΔsdaCB* strains harboring plasmids expressing full-length or truncated SdaC (premature stop codon at E83) in rich medium containing 0.025% (1.4mM) glucose and no serine. For growth curves, three biological replicates are shown as points with their averages connected by lines.

**Fig S4.**
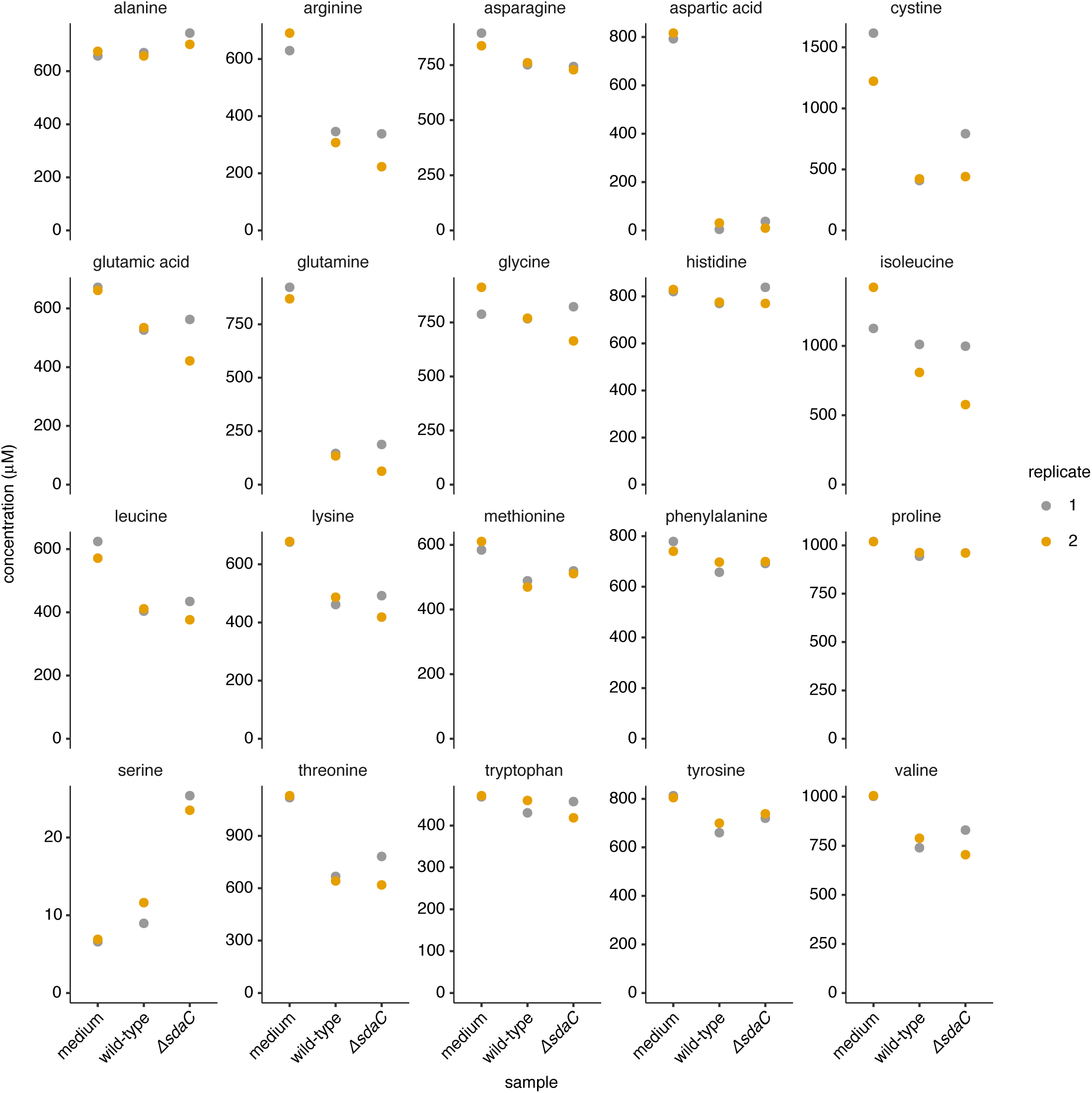
Amino acid concentrations in wild-type vs. *ΔsdaC* culture supernatants. Wild-type and *ΔsdaC* strains were grown to early exponential phase in rich medium containing no serine. Cultures were then passed through a 0.22µm filter and supernatants were stored at –80°C until processing. Amino acid concentrations in culture supernatants and sterile medium were quantified as described in Methods.

## Supplementary Tables

**Table S1.**
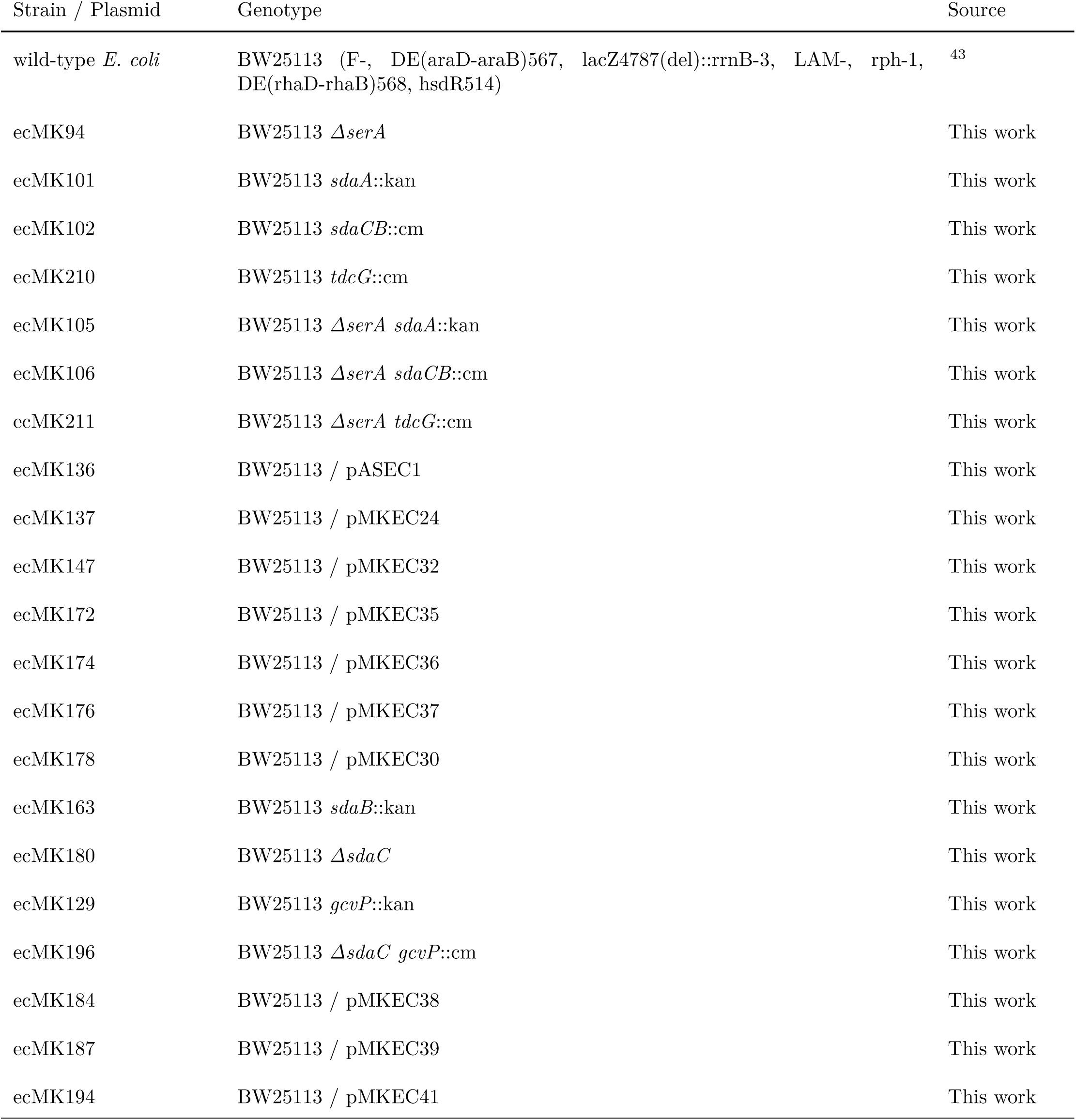
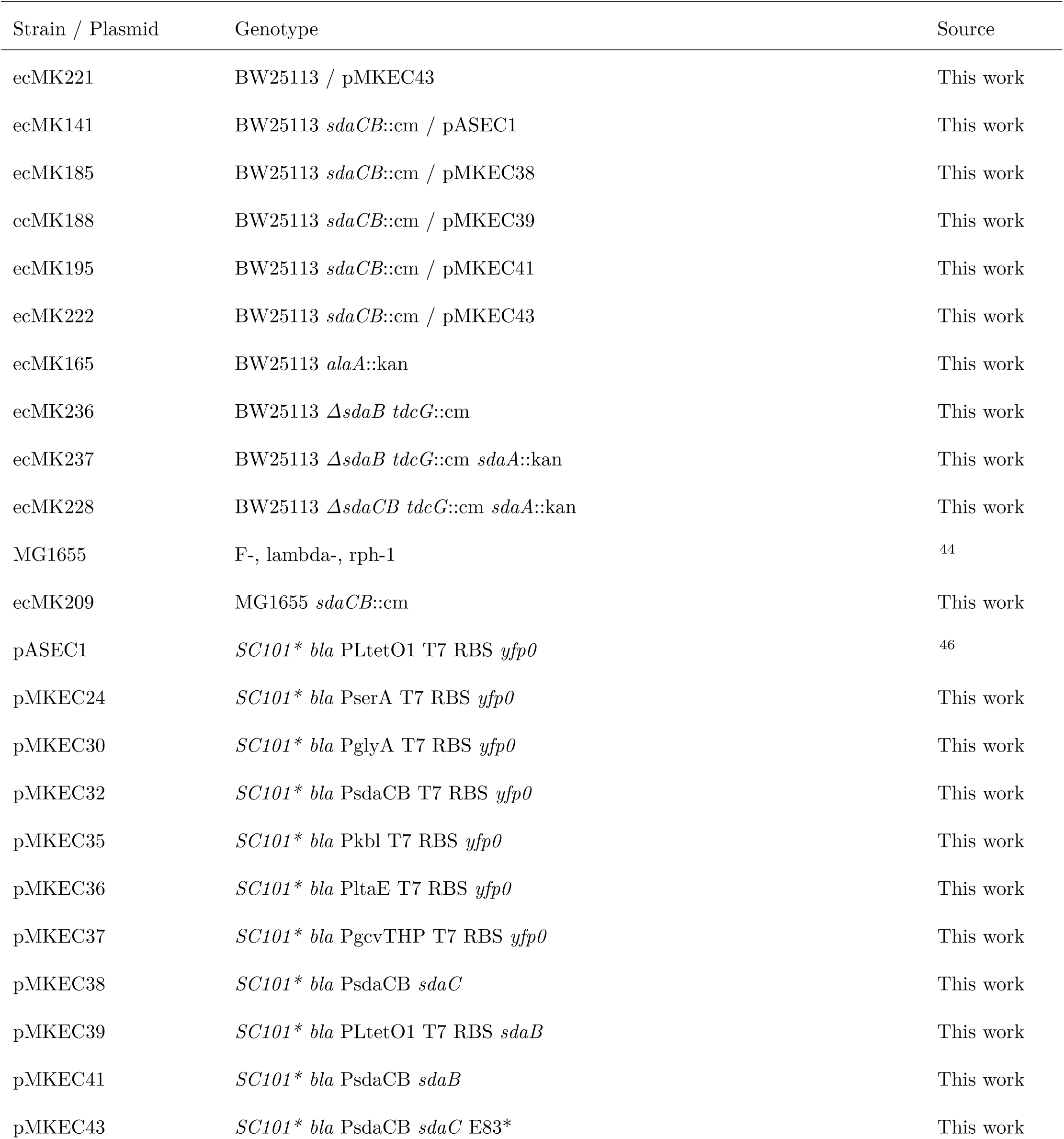
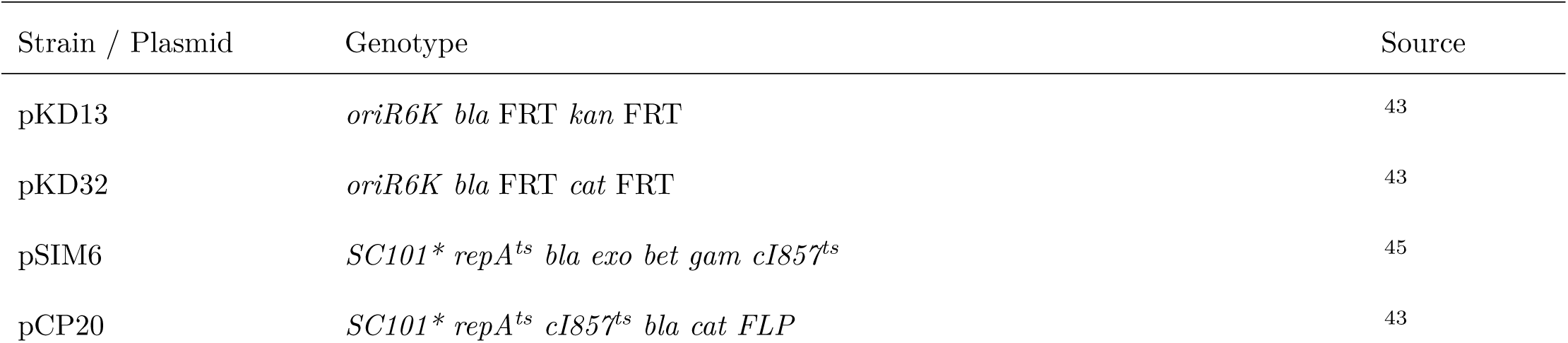
Strains and plasmids used in this study

| Strain / Plasmid | Genotype | Source |
| --- | --- | --- |
| wild-type *E. coli* | BW25113 (F-, DE(araD-araB)567, lacZ4787(del)::rrnB-3, LAM-, rph-1, DE(rhaD-rhaB)568, hsdR514) | <sup>43</sup> |
| ecMK94 | BW25113 $\Delta serA$ | This work |
| ecMK101 | BW25113 *sdaA*::kan | This work |
| ecMK102 | BW25113 *sdaCB*::cm | This work |
| ecMK210 | BW25113 *tdcG*::cm | This work |
| ecMK105 | BW25113 $\Delta serA$ *sdaA*::kan | This work |
| ecMK106 | BW25113 $\Delta serA$ *sdaCB*::cm | This work |
| ecMK211 | BW25113 $\Delta serA$ *tdcG*::cm | This work |
| ecMK136 | BW25113 / pASEC1 | This work |
| ecMK137 | BW25113 / pMKEC24 | This work |
| ecMK147 | BW25113 / pMKEC32 | This work |
| ecMK172 | BW25113 / pMKEC35 | This work |
| ecMK174 | BW25113 / pMKEC36 | This work |
| ecMK176 | BW25113 / pMKEC37 | This work |
| ecMK178 | BW25113 / pMKEC30 | This work |
| ecMK163 | BW25113 *sdaB*::kan | This work |
| ecMK180 | BW25113 $\Delta sdaC$ | This work |
| ecMK129 | BW25113 *gcvP*::kan | This work |
| ecMK196 | BW25113 $\Delta sdaC$ *gcvP*::cm | This work |
| ecMK184 | BW25113 / pMKEC38 | This work |
| ecMK187 | BW25113 / pMKEC39 | This work |
| ecMK194 | BW25113 / pMKEC41 | This work |
| ecMK221 | BW25113 / pMKEC43 | This work |
| ecMK141 | BW25113 *sdaCB::cm* / pASEC1 | This work |
| ecMK185 | BW25113 *sdaCB::cm* / pMKEC38 | This work |
| ecMK188 | BW25113 *sdaCB::cm* / pMKEC39 | This work |
| ecMK195 | BW25113 *sdaCB::cm* / pMKEC41 | This work |
| ecMK222 | BW25113 *sdaCB::cm* / pMKEC43 | This work |
| ecMK165 | BW25113 *alaA::kan* | This work |
| ecMK236 | BW25113 $\Delta sdaB$ *tdcG::cm* | This work |
| ecMK237 | BW25113 $\Delta sdaB$ *tdcG::cm sdaA::kan* | This work |
| ecMK228 | BW25113 $\Delta sdaCB$ *tdcG::cm sdaA::kan* | This work |
| MG1655 | F-, lambda-, rph-1 | 44 |
| ecMK209 | MG1655 *sdaCB::cm* | This work |
| pASEC1 | *SC101\* bla* PLtetO1 T7 RBS *yfp0* | 46 |
| pMKEC24 | *SC101\* bla* PserA T7 RBS *yfp0* | This work |
| pMKEC30 | *SC101\* bla* PglyA T7 RBS *yfp0* | This work |
| pMKEC32 | *SC101\* bla* PsdaCB T7 RBS *yfp0* | This work |
| pMKEC35 | *SC101\* bla* Pkbl T7 RBS *yfp0* | This work |
| pMKEC36 | *SC101\* bla* PltaE T7 RBS *yfp0* | This work |
| pMKEC37 | *SC101\* bla* PgcvTHP T7 RBS *yfp0* | This work |
| pMKEC38 | *SC101\* bla* PsdaCB *sdaC* | This work |
| pMKEC39 | *SC101\* bla* PLtetO1 T7 RBS *sdaB* | This work |
| pMKEC41 | *SC101\* bla* PsdaCB *sdaB* | This work |
| pMKEC43 | *SC101\* bla* PsdaCB *sdaC* E83* | This work |
| pKD13 | *oriR6K bla FRT kan FRT* | 43 |
| pKD32 | *oriR6K bla FRT cat FRT* | 43 |
| pSIM6 | *SC101\* repA<sup>ts</sup> bla exo bet gam cI857<sup>ts</sup>* | 45 |
| pCP20 | *SC101\* repA<sup>ts</sup> cI857<sup>ts</sup> bla cat FLP* | 43 |

**Table S2.**
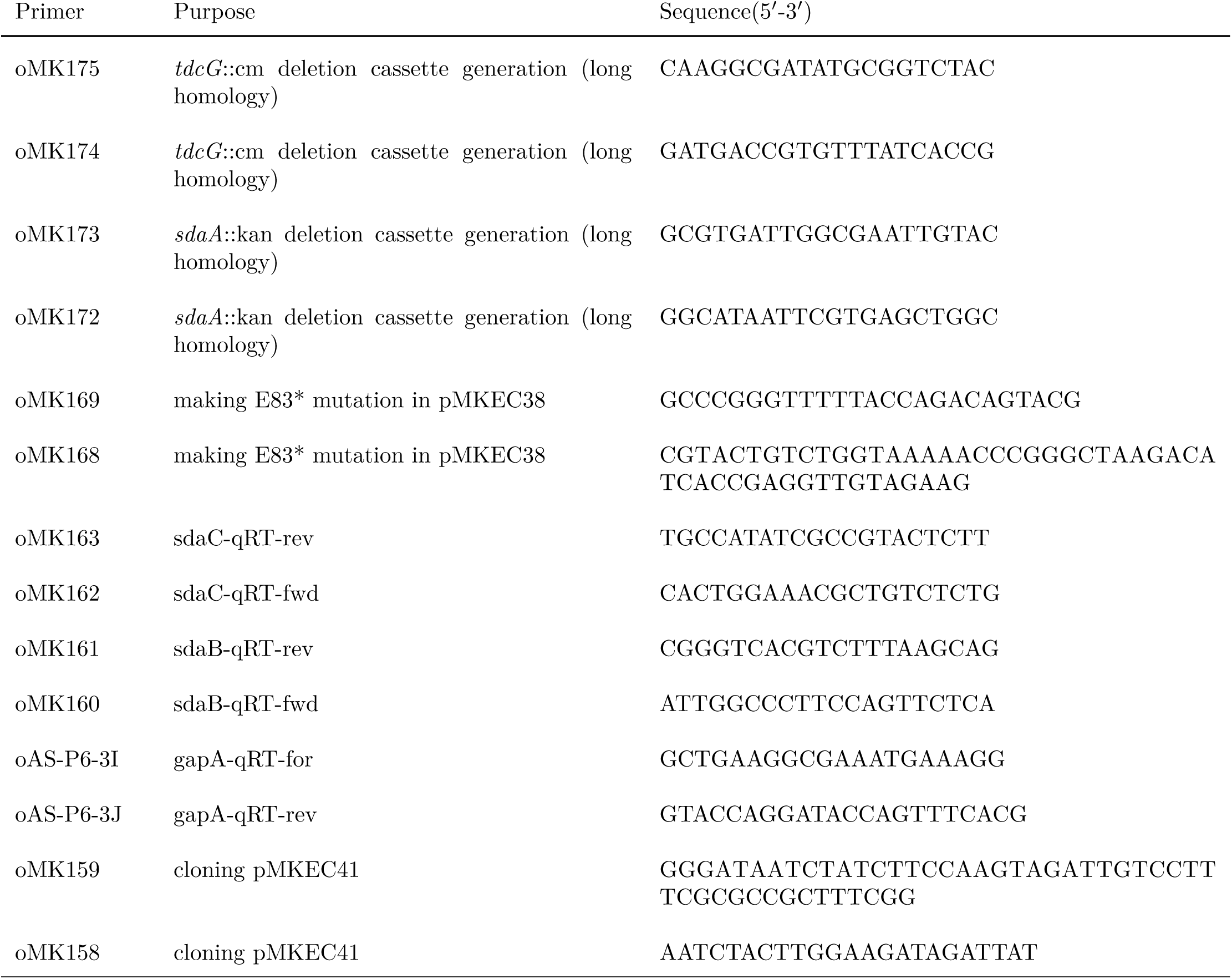
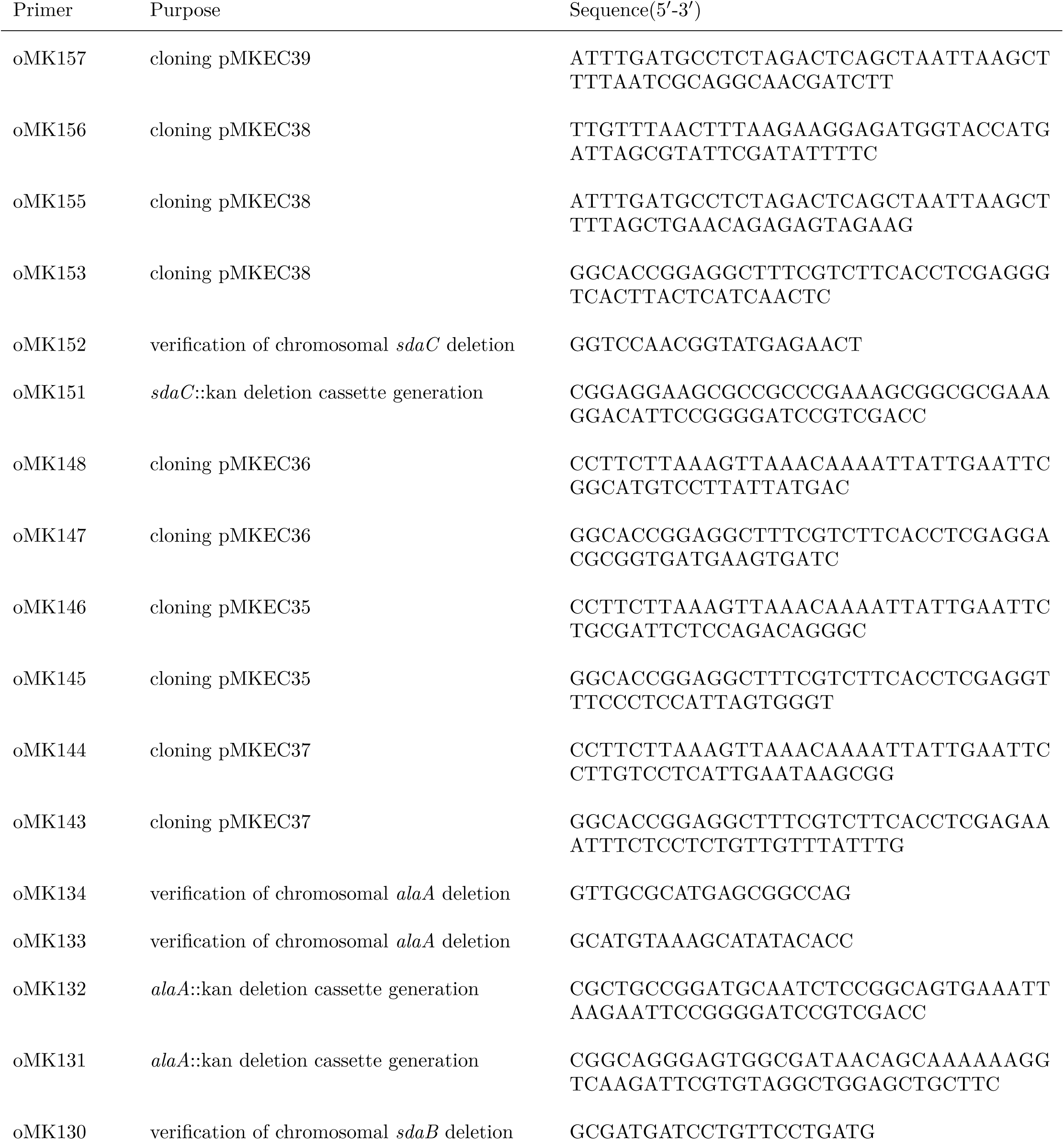
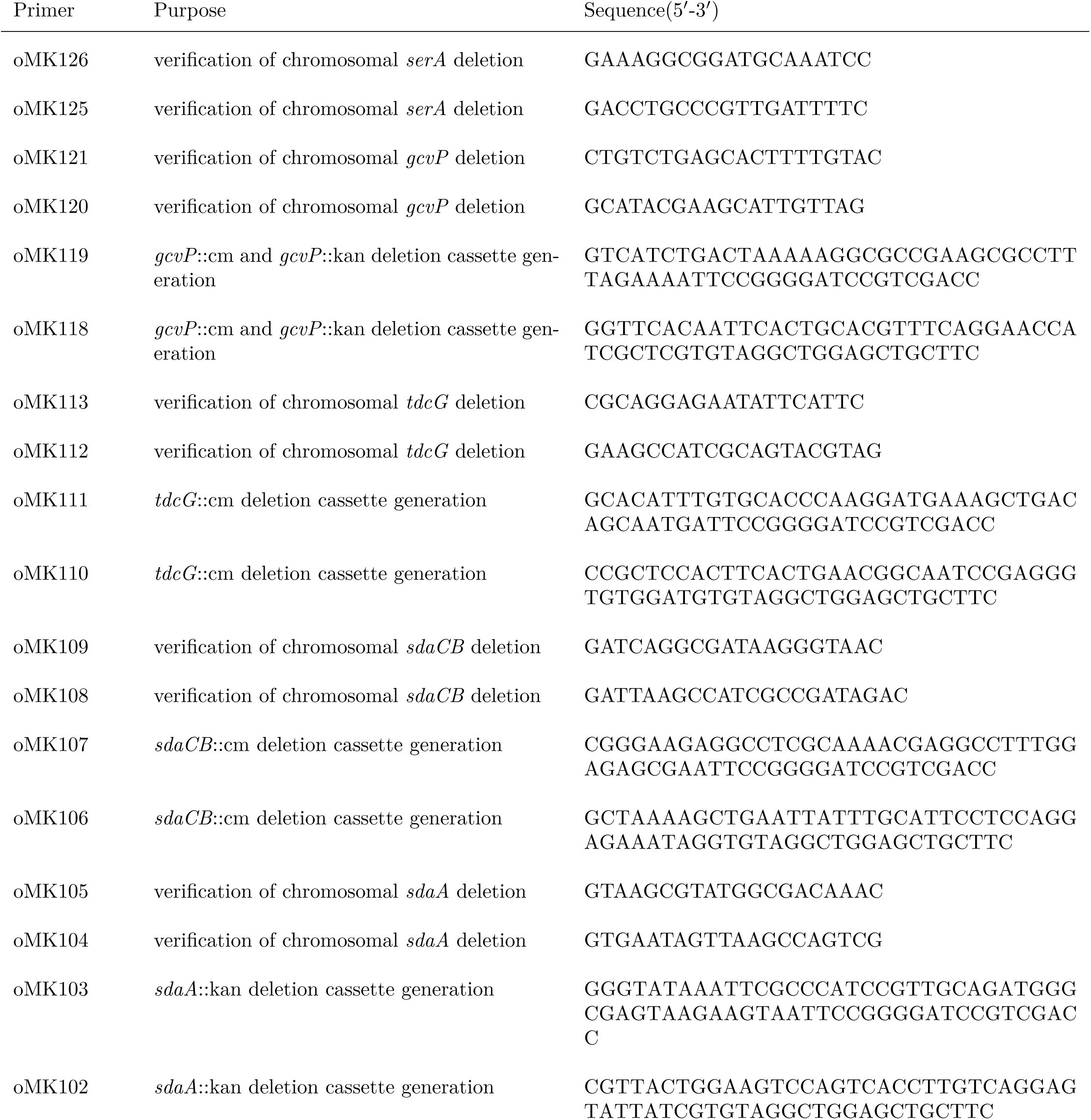
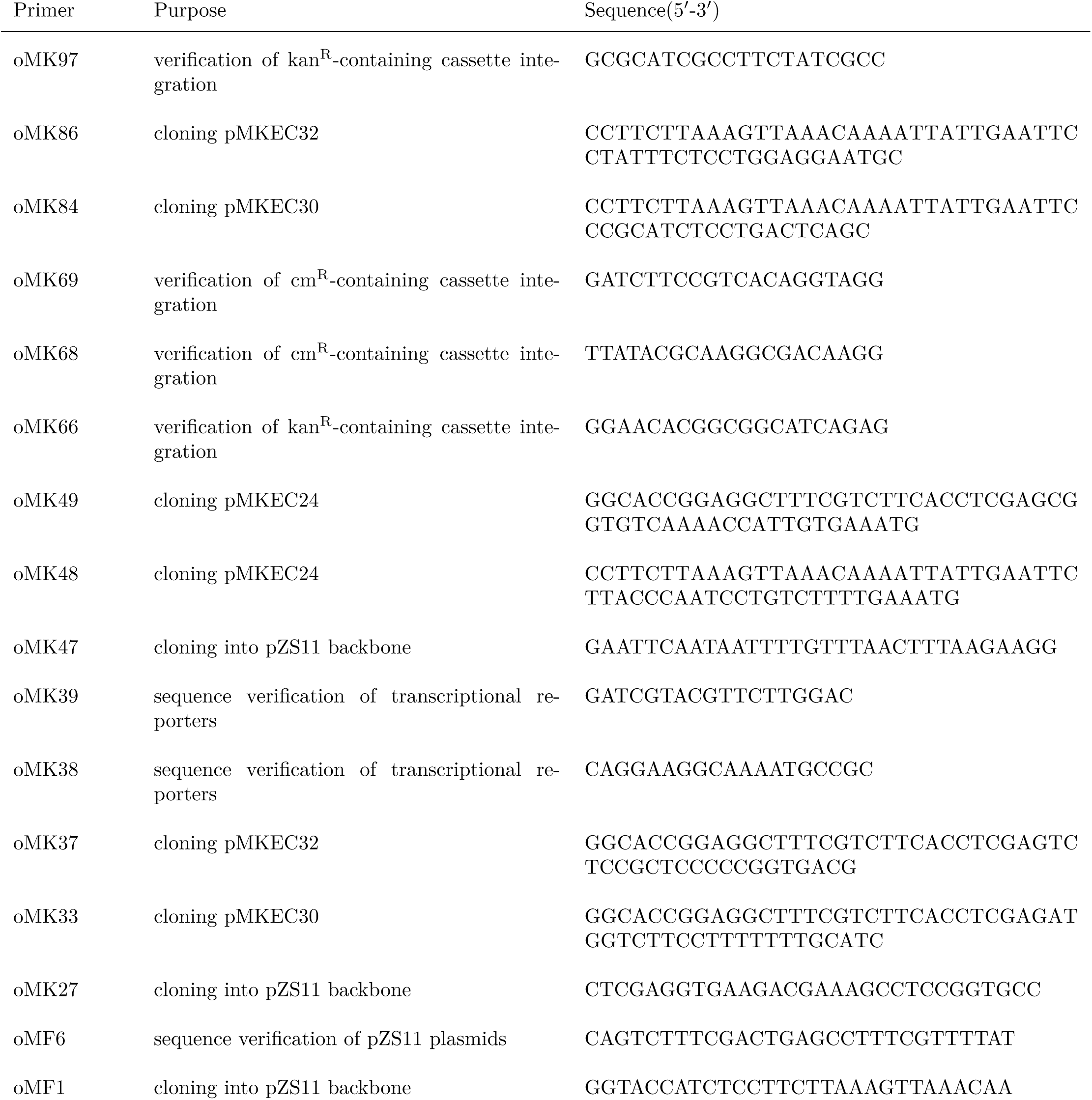
Primers used in this study

